# Expansion of a subset within C2 clade of *Escherichia coli* sequence type 131 (ST131) is driving the increasing rates of Aminoglycoside resistance: a molecular epidemiology report from Iran

**DOI:** 10.1101/855700

**Authors:** Zoya Hojabri, Narges Darabi, Majid Mirmohammadkhani, Romina Hemmati, Zahra saeedi, Kiarash Roustaee, Omid Pajand

## Abstract

The most important lineage of *Escherichia coli*, named sequence type 131 (ST131) is a pandemic clone which drives the increasing rates of antibiotic resistance. While the pervasiveness of ST131 clade C, especially subclades C2 and C1-M27 has been demonstrated in numerous global surveys, no report about the ST131 clades and its virotypes has been published from Iran, so far. So, in this study we investigated and compared the virotypes, antibiotic susceptibility patterns, resistance/virulence determinants and clonality of ST131 clades collected during one-year surveillance study.

Most of isolates belonged to clade C2 (34/76 [44.7%]), with the highest virulence factor (VF) scores and resistance rates. The distinctive profiles of clade C2 virulence genes were revealed by “principle coordinates analysis” (PcoA) test. The distribution of *hlyA*/cnf1virulence genes among clade C2 was not uniform, so that positive strains showed significantly higher rates of resistance markers (*bla*_CTX-M-15_, *bla*_OXA-1_, *aac6Ib*/*Ib-cr* and *aac3IIa*) and ampicillin-sulbactam/gentamicin/tobramycin resistance. Virotype C as the most common virotype (48.7%) was predominant among clade C1 population, while almost all of virotypes E and F [(22/23), 95.6%] strains belonged to clade C2, with the highest VF scores and aminoglycoside resistance rates. “Multi locus variable Number tandem repeats analysis” (MLVA) clustered clades C1 and C2 together, while clades A and B strains were mostly identified as singletons.

Appearance of virotypes E and F among clade C2 strains with higher rates of aminoglycoside resistance/virulence genes content demonstrate the shifting dynamics of this pandemic clone in response to antibiotic selection pressure by establishing the newly-emerged subsets.

## Introduction

Sequence type 131 (ST131), the currently emerged clone of *Escherichia coli* which is disseminated worldwide cause severe hospital-acquired and community-onset infections (*1, 2*). The pervasiveness of ST131 has been reported by many global surveys and increasing prevalence of fluoroquinolone and cephalosporine resistance in *E. coli* population is attributed to this clone (*3*).

Extensive studies of ST131 clone have been started from 10 years ago and it has found to be rapidly expanding across the globe. The first description of this clone was in 2008 (*4, 5*) and has now been found on every continent examined; it is estimated that up to 30% of all ExPEC isolates in some regions belong to ST131 lineage (*6*).

ST131 strains are closely related and appear to have had a common ancestor, so they are often referred as a clone, or clonal group (*7*). However, whole genome sequencing analysis by several studies revealed that ST131 consist of different clades: clade A, B and C (*8*). Generally, the clades A and B which are minor parts of ST131 population, are susceptible to fluoroquinolone and cephalosporine, while clade C (also known as *H*30) represents the largest clade and comprises two sub-clades: C1 (or *H*30R) and C2 (*H*30Rx), both of which are resistant to fluoroquinolone (*8*). The carriage of CTX-M-15 Extended Spectrum β-lactamase (ESBL) gene which is the most widespread β-lactamase among Gram negative bacteria has been shown among C2 subclade (*9*). Despite the highly conserved sequences which are identified in the core genome of ST131, the accessory genome of this clone is much variable and results in differences in virulence gene content and plasmid repertoire (*10*). Considering the virulence genes content, the ST131 clone can be categorized into 12 virotypes which are named from A to F (*11*). While the virotype C is reported as the most common virotype among ST131 clone, the other virotypes have not the equal distribution among ST131 population reported in studies from different continents (*12*).

The detection of ST131 and its clades is important for epidemiological studies. While this being recognized as a pandemic clonal group that threatens public health, ST131 has received less attention in Iran than have other antimicrobial resistant pathogens. Many genotypic tools have been applied to *E. coli*; all of these methods have their advantages and drawbacks (*13*). In a recent study, Multiple-locus Variable Number of Tandem repeats Analysis (MLVA) technique which has been shown as a rapid and highly discriminatory method for typing of *Enterobacteriaceae*, is modified to a single-tube multiplex PCR with standard agarose gel electrophoresis for analysis of the PCR (*14*). So, in this survey firstly we aimed to identify the clades and virotypes of ST131 population and study their clonality using the newly-designed gel based MLVA technique and secondly, determine the differences of virulence and resistance genes content between ST131 clades.

## Results

### Clades determination, O25b/O16 subgroups and virulence genes content

Multiplex PCR for clades determination revealed the C2 clade as the dominant subset [34/76 (44.7%)], followed by C1-M27 [23/76 (30.2%)], C1-nM27 [9/76 (11.8%)], A [8/76 (10.5%)] and B [2/76 (2.6%)]. Strains of clade A were identified as O16 subgroup and fim*H*30 negative, while the remaining 68 isolates including both of the clade B strains belonged to O25b subgroup and harboured the fim*H*30 allele. Seven virulence factors, including *sfa focDE*, *colV*, *tsh*, *vat*, *papGIII*, *cdtB* and *neuCK* were not detected among study isolates. The 27 virulence markers were detected at least once, with the lowest rate of 1.3% (*ompT*, *hlyF*, *iss*) to the highest rate of 100% (*usp, yfcV, fyuA, chuA*). Except for two C1 sub-clades (C1-M27, C1-nM27), the other three clades were considerably different in VF content, with the lowest VF score of clade A (median: 11) to clade C2 with the highest VF score (median: 15).

Among the C1 sub-clades, capsular types *kpsMTII* and *k5* were significantly detected among C1- M27, while *papACEFGII*, *hra*, *hlyA* and *cnf1* were negatively associated with both of the C1 sub-clades. Seven virulence markers (*hra*, *cnf1*, *hlyA*, *papA*, *papGII*, *papC* and *papEF*) were significantly associated with clade C2. Furthermore, positive association was found between the carriage of *hlyA* and *pap*ACEFGII, *iha*, *hra* and *cnf1* within this clade.

Principal coordinate analysis based on the 27 virulence determinants revealed that virulence profiles of clade C2 were distinctive and differentiated them from other clades. Plotted coordinate 1-coordinate 2 plane showed that the variance of 65.29 was captured and clade C2 isolates were clustered in the lower right quadrant, clearly separated them from other clades (Figure 1). The differences of aggregate virulence profile were explored using univariate analysis. Clade C2 strains showed higher aggregate virulence score (median) than other clades. All isolates fulfilled molecular criteria for UPEC, while 72 (94.7%) strains were identified as ExPEC. Table 1 shows the prevalence of virulence genes among different clades.

**Figure 1.**
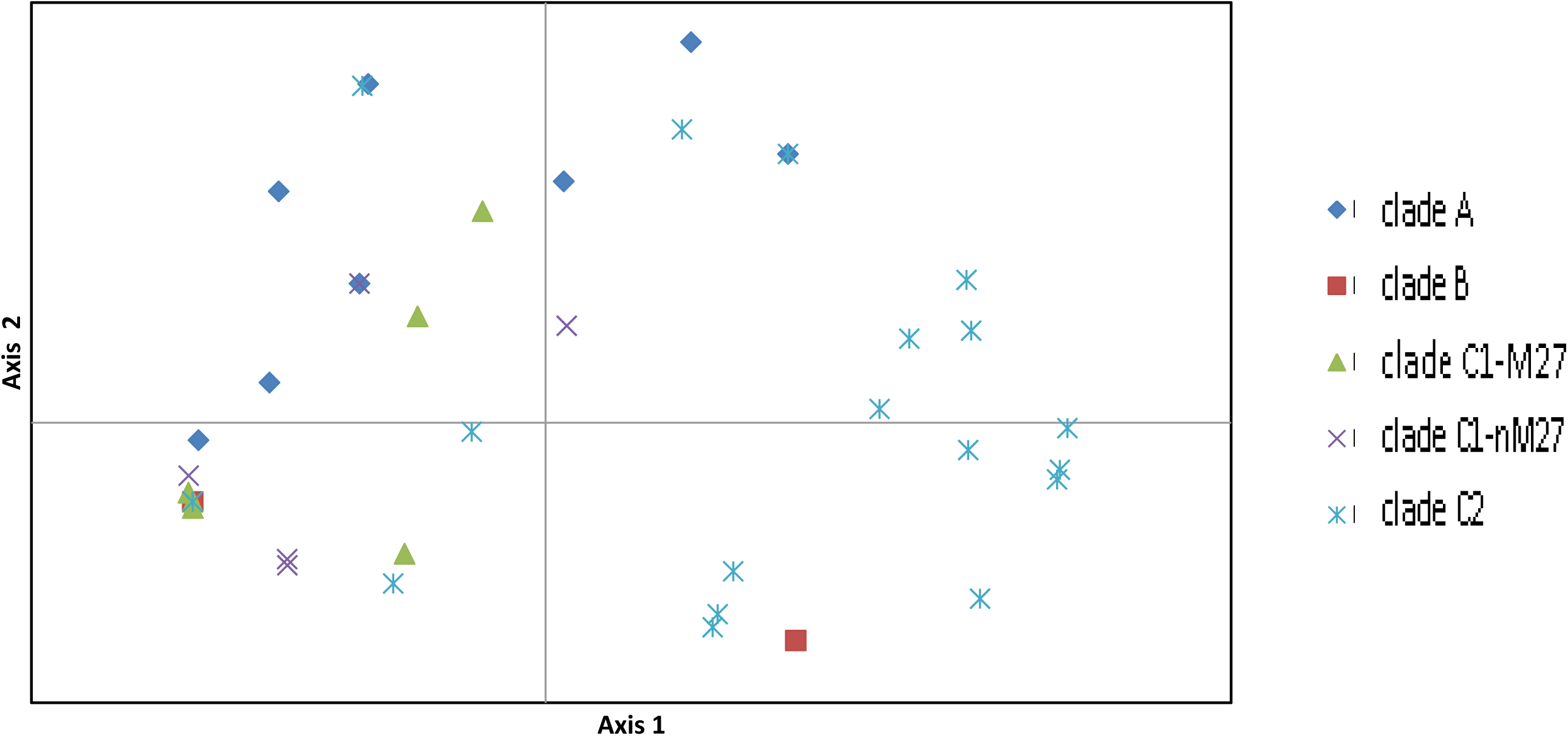
Principal coordinate analysis (PCoA) of virulence gene profiles among 76 ST131 isolates. The PCoA was based on results for all 27 virulence genes studied. Each isolate is plotted based on its values for PCoA coordinates 1 (*x* axis) and 2 (*y* axis), which collectively capture 65.29% of total variance in the data set.

**Table 1.**
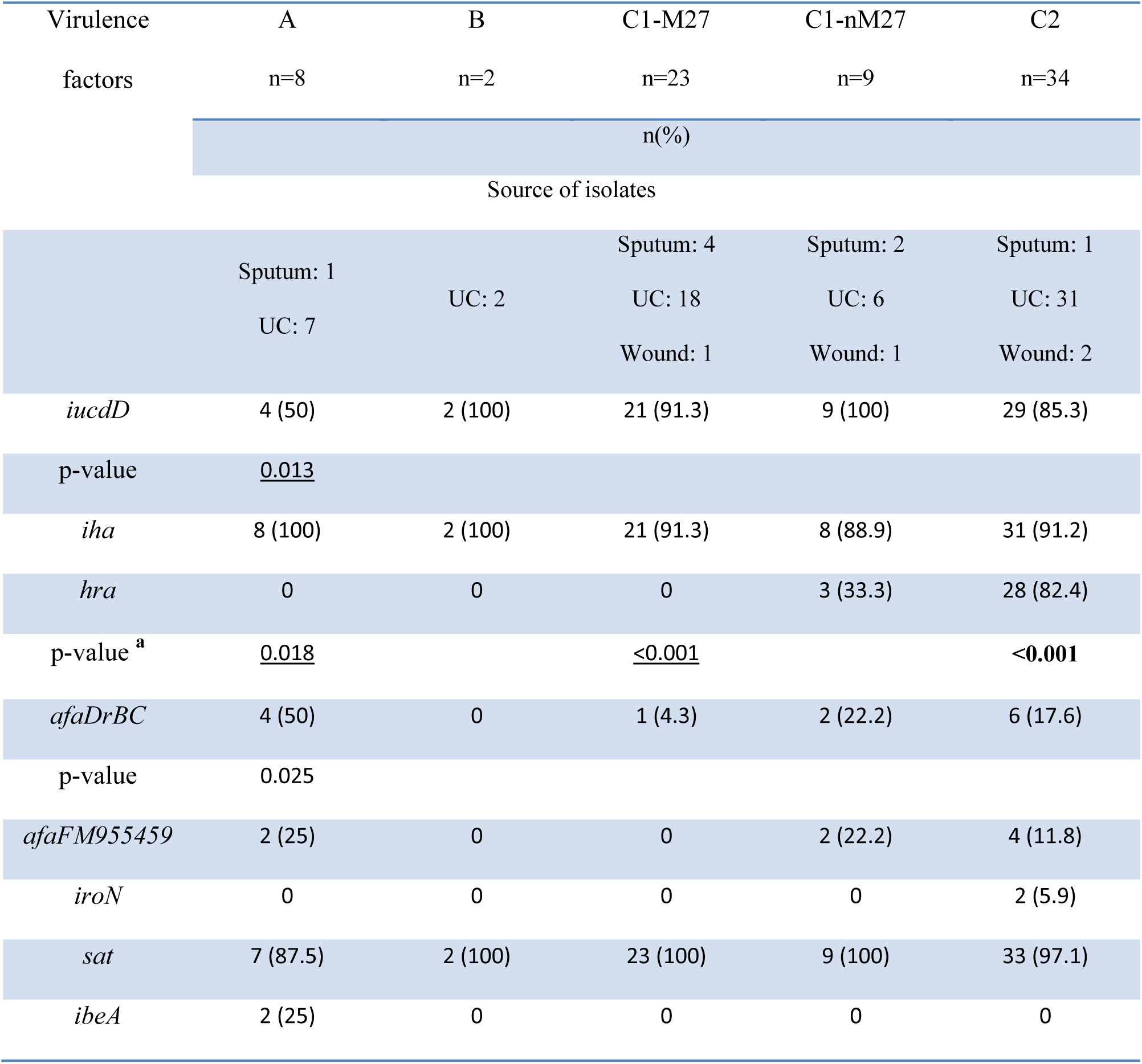
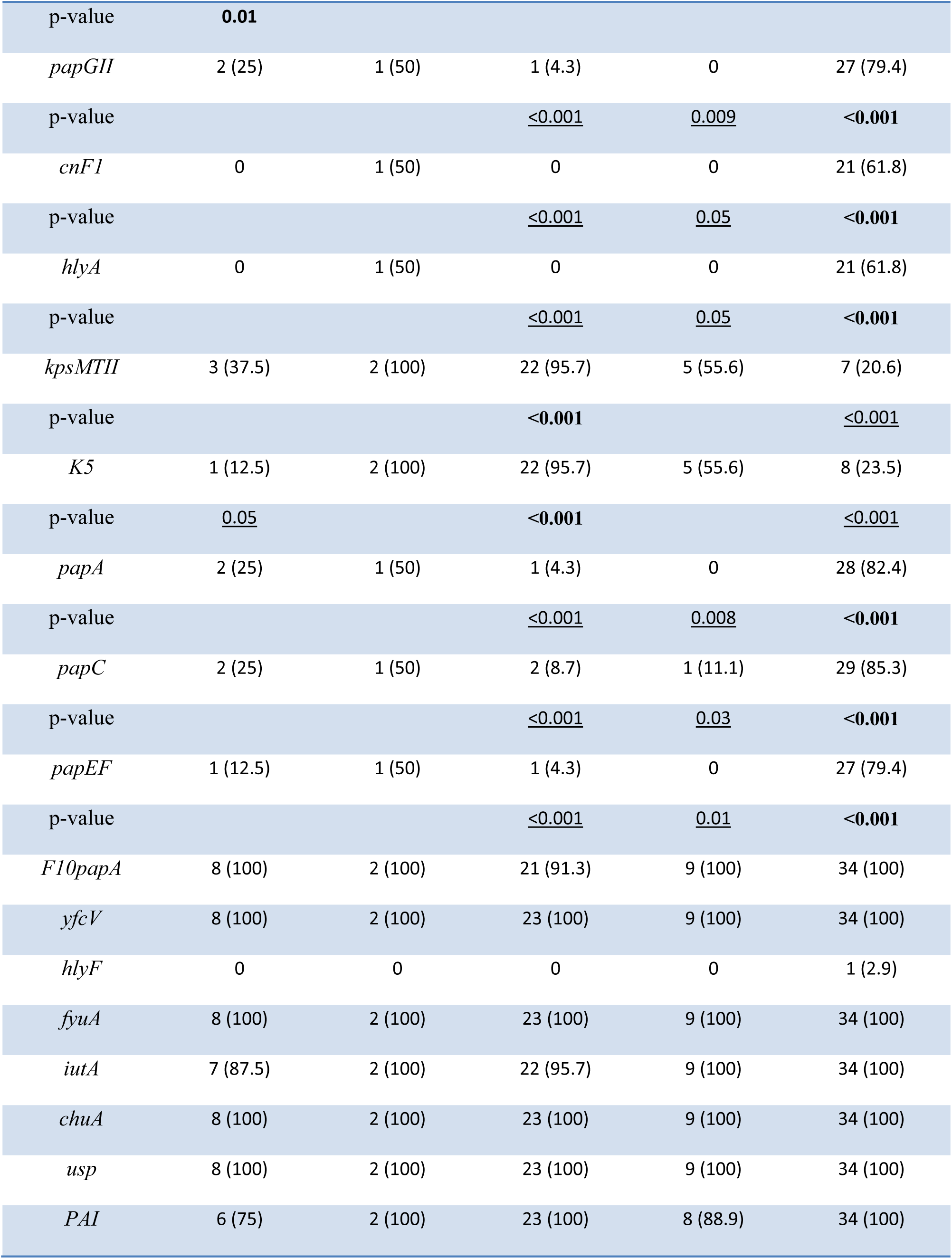
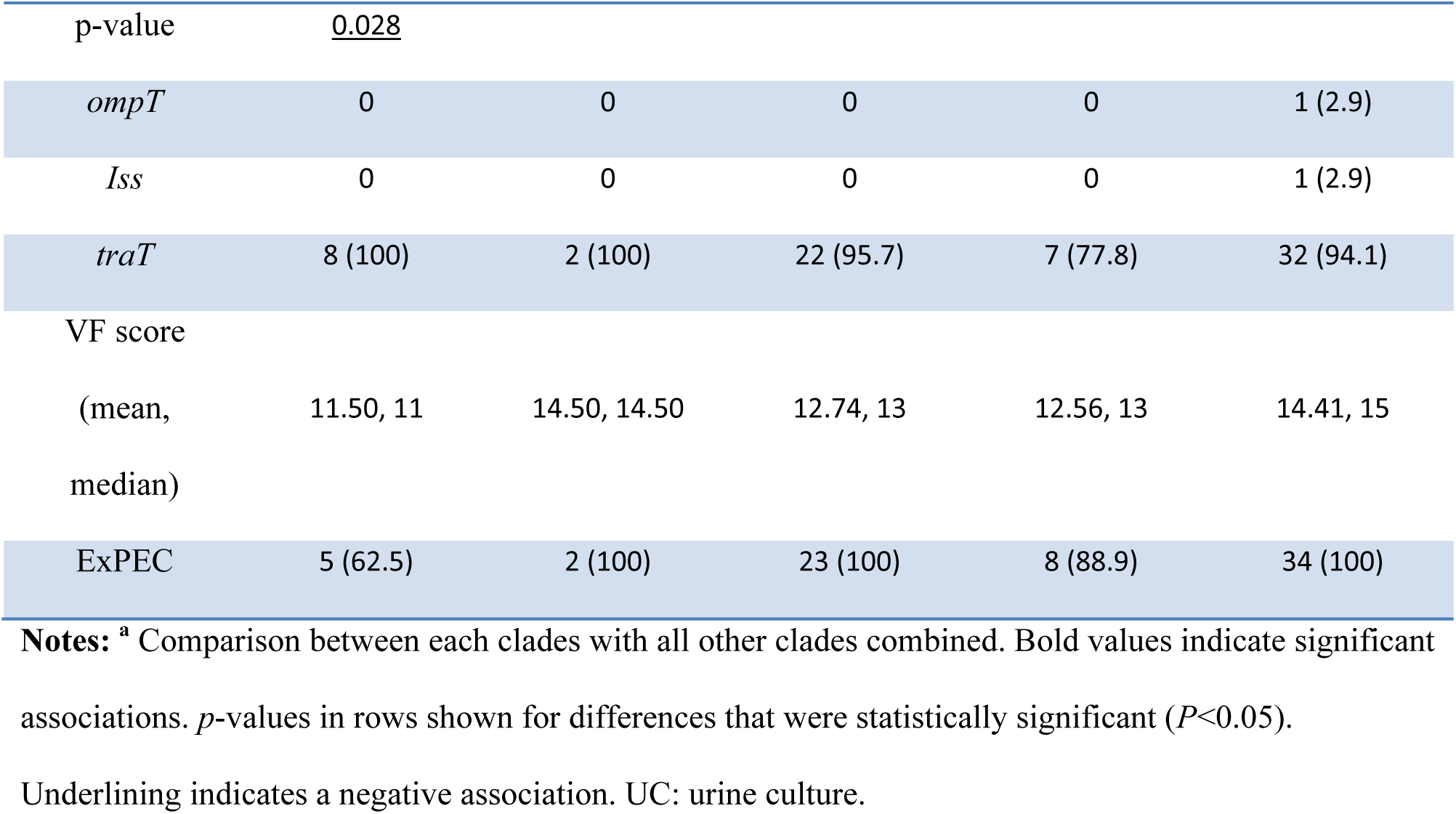
Prevalence of virulence markers among ST131 clades.

### Clades and resistance profiles/resistance determinants

Clade A was significantly associated with susceptibility to fluoroquinolone and tobramycin, while the difference was borderline for the latter antibiotic. The two C1 sub-clades, showed different patterns of antibiotic susceptibility profile. The C1-M27 was found significantly associated with susceptibility to ampicillin/sulbactam, amoxicillin/clavulanate, gentamicin and tobramycin, and was resistant to fluoroquinolone, however, no such an association was found for the C1-nM27 subclade. The resistance rates were increased from clade A with median of 3 to clade C2 (median: 5). Clade C2 isolates showed significantly higher rates of resistance against gentamicin, amikacin, tobramycin, amoxicillin-clavulanate, ampicillin-sulbactam, nitrofurantoin, and fluoroquinolone. Of the resistance markers studied, *aac6-Ib/Ib-cr*, *bla*_OXA-1_ and CTX-M-G1 were associated with clade C2 strains, while this clade was conspicuous for the low prevalence of *bla*_TEM-_ (*P* = 0.01) (Table 2).

**Table 2.**
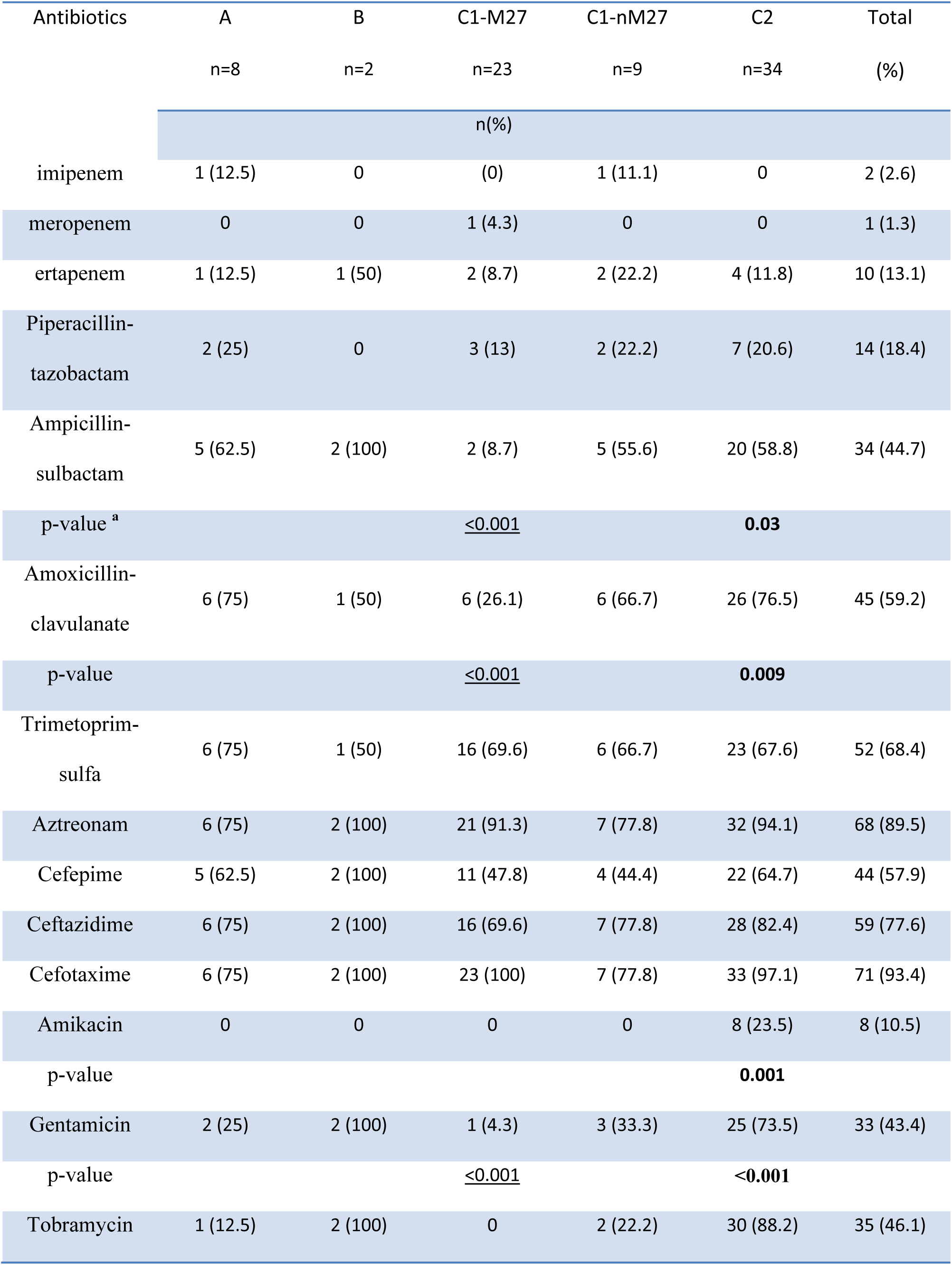
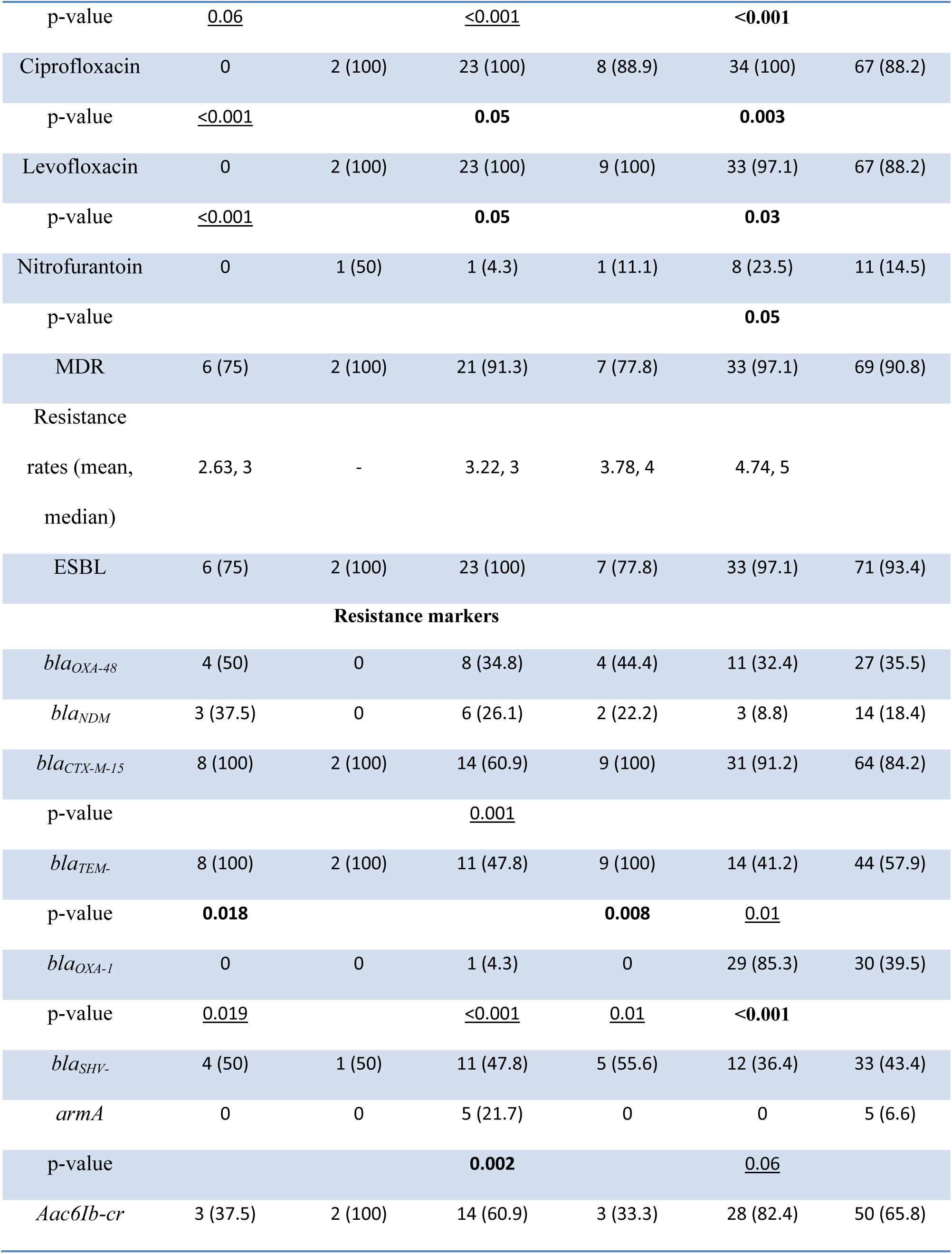
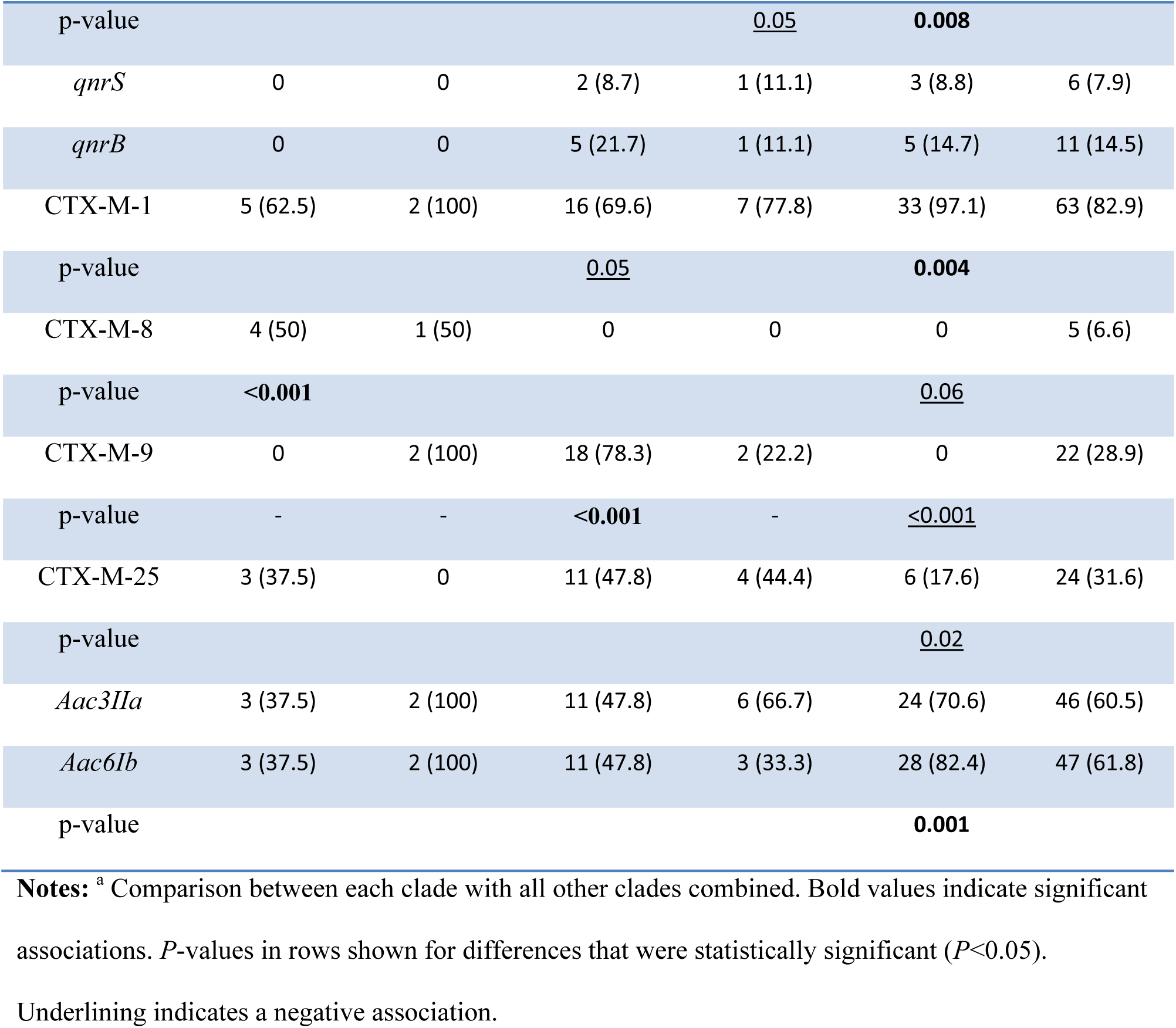
Antibiotic resistance rates and prevalence of resistance markers among different clades of ST131.

Closer examination of the C2 clade showed that *hlyA* gene wasn’t uniformly distributed among this clade and strains carrying this virulence marker were significantly positive for *aac6Ib* (*P* = 0.001), *aac6Ib-cr* (*P* = 0.02), *bla*_CTX-M-15_ (*P* = 0.048), *bla*_OXA-1_ (*P* = 0.005) and *aac3IIa* (*P* = 0.002). The carriage of *hlyA* among C2 clade was coincided with resistance phenotype to tobramycin (*P* = 0.015) and gentamicin (*P* < 0.001). The CTX-M-G1 and CTX-M-G9 were significantly more common in C2 and non-C2 clades, respectively. The *bla*_CTX-M-15_ was detected in more than 90% of all clades, except for C1-M27 which had negative association with this resistance marker. The *bla*_CTX-M-15_ positive isolates exhibited a significantly higher prevalence of resistance to ampicillin/sulbactam (*P* = 0.009), tobramycin (*P* = 0.004) and gentamicin (*P* = 0.01).

Considering the *bla*_OXA-1_, there was a strong association between the carriage of this element and resistance phenotype to amikacin, gentamicin, tobramycin, aztreonam, amoxicillin/clavulanate, ampicillin/sulbactam and fluoroquinolone. Also, strains carrying the *bla*_OXA-1_ were significantly positive for the *bla*_CTX-M-15_ and *aac6Ib*/*Ib-cr*.

### Virotyping of clades and resistance markers

All except four ST131 strains divided in to virotypes A to F. The remaining four strains (no. 502 [O16] and 418A, 672, 81 [O25b]) showed unknown virulence genes patterns and couldn’t be categorized into virotypes. Virotype C was the most common, represented by 37/76 (48.7%) isolates and predominantly associated with clade C1 (29/37), including subclade C1-M27 [22 strains within virotype C, (59.5%)] and C1-nM27 [7 strains within virotype C, (18.9%)]. In contrast, all but one of the virotype E strains belonged to clade C2 [19 strains within virotype E (95%)] (Table 3).

**Table 3.**
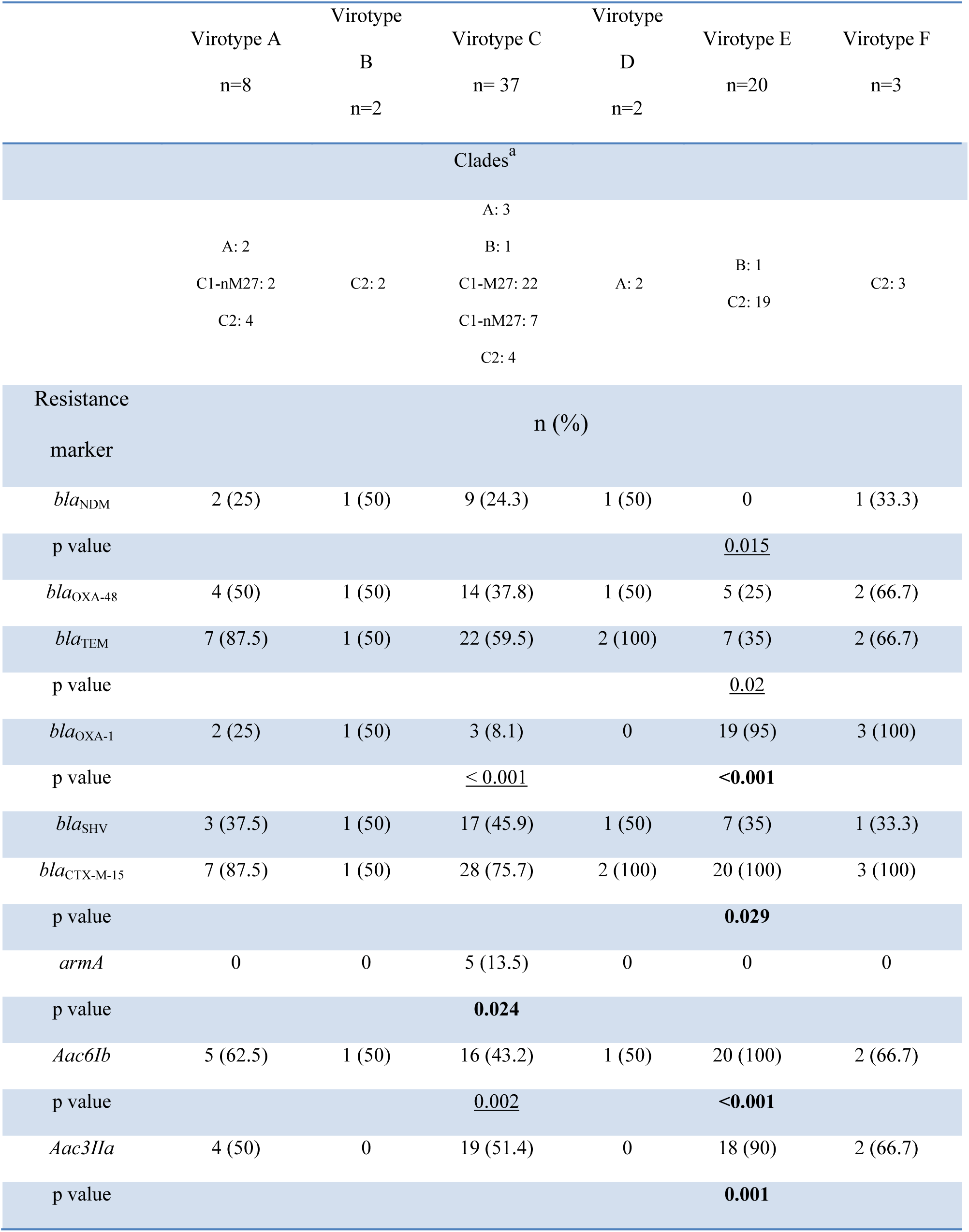
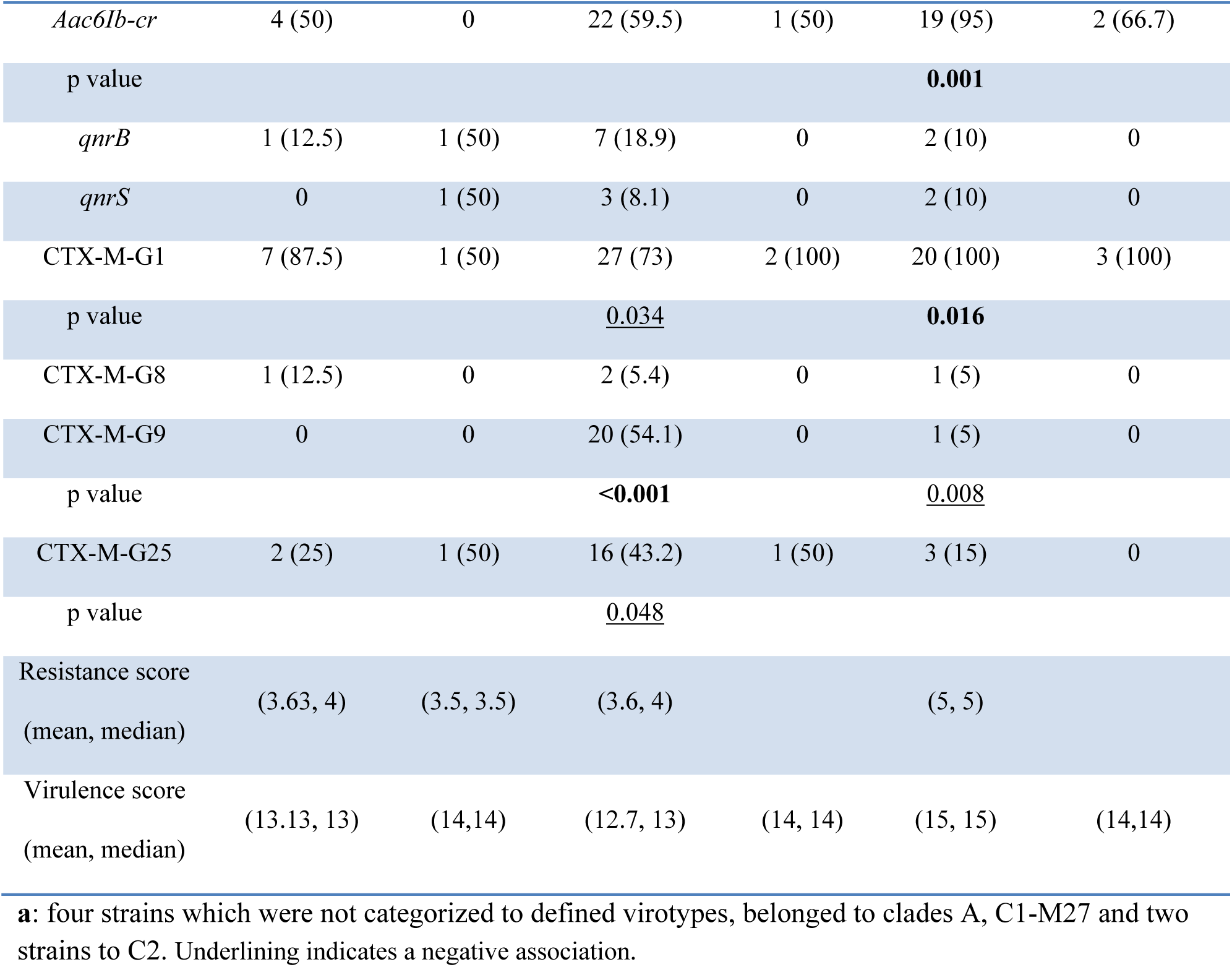
Virotyping of clades and prevalence of resistance genes among different virotypes.

The associations among all 27 detected virulence genes were identified using cluster analysis. The isolates were divided according to 100% similarity of virulence genes content. The largest cluster included 37 strains with a unique set of six virulence genes (*sat*, *chuA*, *fyuA*, *yfcV*, *usp* and *iutA*) that corresponded to virotype C. These isolates harbored mostly *bla*_CTX-M-15_ (28/37; 76.7%), or were *bla*_OXA-1_ negative (31/34; 91.8%), and mainly represented the C1 clade (29/37; [78.4%], including 22 C1-M27 and seven C1-nM27 strains).

The second largest cluster, which included 19 clade C2 strains, contained mostly *bla*_OXA-1_ (100%), *bla*_CTX-M-15_ (100%), *aac3IIa* (89.5%) and *aac6Ib*(100%) */aac6Ib-cr* (94.7%) positive strains with a set of 12 virulence markers (*papACEFGII*, *F10papA*, *sat*, *cnf1*, *hlyA*, *chuA*, *fyuA*, *yfcV*, *iutA*, *iha*, *traT*, *usp* and *malX*) that corresponded to virotype E. As expected, virotype E strains were significantly associated with resistance phenotypes to ampicillin/sulbactam (*P* = 0.04), gentamicin (*P* < 0.001), tobramycin (*P* < 0.001) and amikacin (*P* = 0.003).

The highest VF score was detected among virotype E (mean and median: 15 for both), followed by virotypes B/F (mean and median: 14 for both virotypes), virotype A (mean: 13.13, median: and virotype C (mean: 12.70, median: 13). ExPEC criterion was predominated among *bla*_CTX-M-15_ carrying strains (60/64, 93.7%).

### MLVA typing of ST131 strains

Based on the MLVA typing, 18 profiles were identified among 76 ST131 strains, including two to 11 isolates in each cluster and 5 strains were found singletons. Considering the clades, clade A which included eight O16 strains was divided in to two clusters and three singletons. Two clade B strains were also identified as singletons. The MLVA generated dendogram grouped clades C1 and C2 in the same clusters, in comparison with clades A and B which had different and distinct banding patterns and were not clustered together (Figure 2).

**Figure 2.**
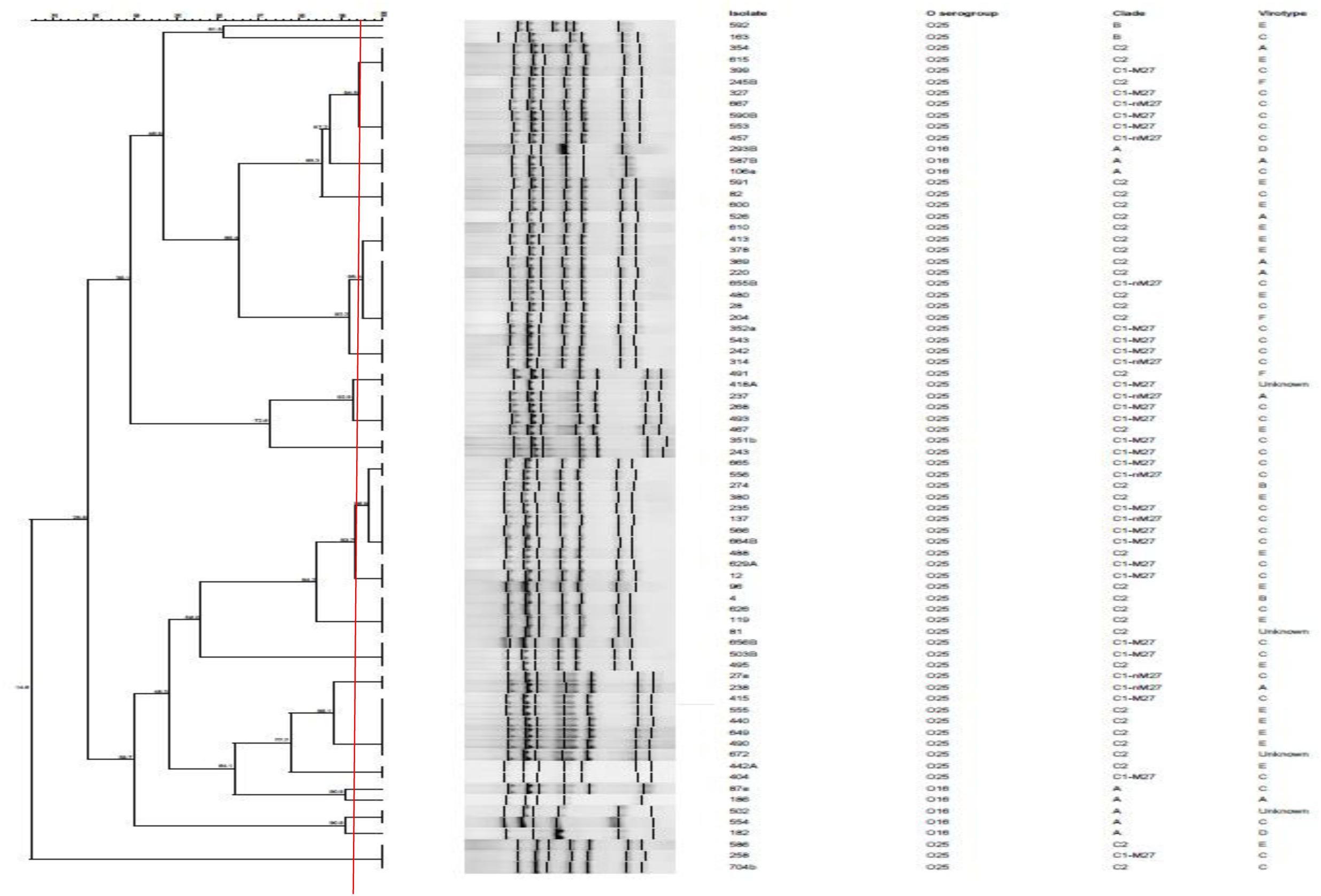
MLVA-based dendrogram of 76 ST131 *E. coli* isolates. The dendrogram was constructed by BioNumerics software using UPGMA and the Dice algorithm based on band profile on gel electrophoresis. The red line corresponds to the 95% similarity cut-off. O25b/O16 subgroups, clades and virotypes are shown.

## Discussion

The worldwide spread of the ST131 clone of *E*. *coli* is a notable example of the rapid expansion of a successful pathogen (*3*). Multiple researchers examining ST131 strains from different countries have been reported that C1 and C2 clades are the dominant subsets among ST131 population and drive the global success of this clone (*15*).

As far as we know, this is the first study in Iran to investigate and compare the prevalence and genotypes of ST131 sub-clades. Here, we found that the C2 clade of ST131 was responsible for most of the ST131 infections. All clades were detected among study population and clades C1 and C2 strains had similar MLVA banding patterns and grouped in a cluster, while clades A and B were different and mainly detected as singletons. Interestingly, virotype and clade patterns had consistency, in which the most common virotypes including virotypes C and E had comprised the clades C1 and C2 strains, respectively. A subpopulation among clade C2 lineage was detected which showed higher carriage rates of resistance/virulence markers and were particularly resistant to aminoglycosides, confirming the importance of emerged subsets within this clone.

In our survey, ST131 strains obtained from different extraintestinal specimens represented the internationally important clades, mainly C1 and C2, suggesting the successful spread of these lineages within studied tertiary hospital. As it is clear, the resistance is increased from C1-M27 to C1-nM27 and C2 strains. While C1-M27 strains were found as the second most prevalent subset among studied ST131, this clade was significantly susceptible to first line antibiotics, including ampicillin/sulbactam, amoxicillin/clavulanate, gentamicin and also tobramycin. This finding indicates that we need to investigate factors other than antibiotic selection pressure to explain this paradox. In contrast to C1-M27, clade C2 strains found resistant to almost all of antibiotics. This observation was in concordance with increasing rates of resistance genes content from clade C1 to C2, particularly CTX-M-G1, *bla*_CTX-M-15_, *bla*_OXA-1_, *aac6Ib/Ib-cr* and *aac3IIa*, which are the most important determinants in developing resistant phenotypes among gram negative pathogens.

Despite the lower prevalence of clades A and B as compared to clade C, they showed important features. First, higher rates of the two most widespread carbapenemase genes including *bla*_OXA-48_ and *bla*_NDM_ along with CTX-M-G8 ESBL were found among clade A strains. Moreover, resistance rates against piperacillin/tazobactam, amoxicillin/clavulanate, trimethoprim/sulfametoxazole in clade A, and ertapenem, ampicillin/sulbactam, cephalosporins, tobramycin, gentamicin, nitrofurantoin and interestingly fluoroquinolone in clade B were higher than clade C, while the differences were not statistically significant. Furthermore, the only fosfomycin resistant strain belonged to clade B (data not shown). The predominance of *bla*_OXA-48_ among mostly fluoroquinolone susceptible clade A strains has been reported in a recent survey from England (*16*). So, it seems that ST131 clone has the ability to compensate the lower prevalence of clades A and B which are usually found as minority subsets by the propensity to accumulate resistance determinants and offset this deficiency.

Virotype C is considered to be the most widely distributed virotype among ST131, occurring in all ST131 clades (*17*). Here, virotype C was predominant among clade C1, specifically C1-M27 subclade, as also reported recently from South-West England and Europe (*17, 18*). In contrast to several European studies which reported the virotype A as the dominated virotype among C2 clade (*12*), in our study virotypes E and F were found as the most common virotypes, representative of > 60% of the C2 clade strains (22 out of 34 strains). Clade C2 showed a heterogeneous population based on the carriage of *hlyA* virulence marker, as positive strains had higher rates of resistance genes content and consequently higher resistance phenotype to tobramycin and gentamicin. Interestingly, almost all of this C2 subclade (19 *hlyA*+/21 *hlyA*+) was identified as virotype E. In a recently published data from Southeast Asia, a subpopulation among C2 clade is reported which has been named Southeast Asia-C2 (SEA-C2) lineage (*19*).

The main features attributed to this subset were the higher carriage rates of some virulence genes mainly *hly*ABCD operon, *cnf1* and *tia/hek*, and also harbouring a conserved plasmid which carried *aac3IIa*, *aac6Ib/Ib-cr*, *bla*_CTX-M-15_, *bla*_OXA-1_ and *tetA*. The strains were very closely related by genome sequence analysis. These findings in conjunction with our data suggest that virotypes E and F are more prevalent than other virotypes among C2 clade strains originated from Asia, constituting a distinct subset among C2 clade population.

With respect to virulence factors, again, high similarity was observed among C2-subclade which were identified as virotype E and probably accounts for some of the virulence factors being overrepresented (as supported by cluster analysis). Some of these genes are commonly present in many ExPEC, particularly UPEC. Notably, the *pap* gene locus which is specifically play important role in causing pyelonephritis (*20*). The second one is hemolysin coding operon, including *hlyA* and *hlyF*, which are shown to harboured mainly by the other major sequence types, including ST73, ST95 and ST127 (*21*). The expression of this virulence marker in the context of urinary tract infections causes the inflammatory cell death of bladder epithelial cells (*22*). The *cnf1* and *hra* are the other two most important virulence factors which have important roles in creating cytopathic effect in infected epithelial cells and adherence, respectively (*23*). Overall, while the strains harboring these virulence factors might have priority to other ST131 population, their overrepresentation in the virotype E is probably due to their close genetic relatedness among C2 clade.

In the present study, we used a recently new-designed MLVA technique in which the analysis of PCR products was based on gel electrophoresis (*14*). Using this molecular typing method, ST131 strains were categorized into 18 profiles. However, we couldn’t establish an association between the generated clusters and clades or virotypes. As detected, the two less common clades A and B, could be differentiated from each other and also from clade C strains, while two clade C subsets were grouped in common clusters. In addition, the differentiation of *hlyA* carrying strains as compared to negative ones couldn’t be distinguished by this method. Indeed, the main reasons we chose the MLVA rather than other gel electrophoresis-based typing methods were its high reproducibility and also superior discriminatory power relative to DiversiLab REP-PCR which were claimed by Caméléna et al (*14*). So, the distinctiveness of our C2-subclade should be certainly confirmed by other robust methods such as pulsed field gel electrophoresis (PFGE).

In conclusion, our study is notable for examining a one-year collection of ST131 strains in a geographical region from which no data has been previously published. This will help us to characterize the local microbiology in this region and the resistance profile among ST131 strains. Our study identified two clade B strains which were O25b/*H*30+ and ciprofloxacin resistant, despite what we had knew about their characteristics so far. We have found that study ST131 strains aren’t a uniform population, and clade C2 like in other regions is driving both the higher virulence and resistance among this high risk clone. More focus on this clade identified a subset of strains which showed virotype E pattern and their resistance phenotypes and resistance/virulence genes repertoire were different from the other C2 clade strains. In concordance with a recent study from Singapore (*19*), our data show a local trend within the C2 clade in which the generated subset has a major advantage over other ST131 population in the context of resistance/virulence genes content and clonality. So, ongoing monitoring of the dynamics of ST131 local transmission is required to understand the reasons of new virotypes emergence within the region.

## Material and methods

### Strains

In this one-year cross-sectional study, 76 non-duplicate phylogroup B2 ST131 isolates were cultured from patients with extraintestinal infections admitted to Kosar university hospital of Semnan, Iran. The clinical samples were collected as part of standard care for admitted patients. Isolates were cultured from different specimens including urine, blood, wound and respiratory samples. These ST131 strains were identified from a collection of 338 *E. coli* isolates based on the *gyrB*/*mdh* single nucleotide polymorphism (SNP) multiplex PCR (*24*). The O25b/O16 subgroups were determined as described earlier (*25*). Allele specific primers for allele 30 of fim*H*30 corresponding with the main fluoroquinolone resistance associated subset within this clone were used to identification of *H*30 subclone (*26*).

### Clades determination and MLVA-typing

For determination of ST131 clades, the multiplex PCR using seven pairs of primers was used as described by Matsumura et al (*27*). Each clade was identified based on the expected amplicons. Amplification was performed using the Ready to use Master Mix (Tempase 2X master mix, Amplicon, Denmark) and recommended concentrations of primers (*27*). For MLVA typing, seven VNTR regions were amplified by a single-tube multiplex PCR in a 25 µl final volume using the Tempase 2X ready to use Master Mix (Amplicon, Denmark) (14). The PCR run conditions and primers concentrations were the same as described by Caméléna et al. The PCR products were loaded to 3% agarose gel and electrophoresed during 75 minute at 100 V. the TIFF image file of gel electrophoresis was saved and gel images were loaded in to a Bionumerics database. MLVA patterns were compared with a tolerance parameter of 1% and an optimization parameter of 0.5%. Using Bionumerics software, the generated dendogram was analyzed by UPGMA and pairwise Dice similarity coefficient. Identical profiles were those that displayed the same banding pattern (corresponding to >95% similarity on dendograms) (*14*).

### Virulence factors and virotype determination

The presence of 34 putative virulence markers was assessed by multiplex PCR (*28, 29*). The carriage of ≥ 2 virulence genes, including *afa drBC*, *papAH* or *papC*, *sfa focDE*, *kpsMTII* and *iutA* was considered as the criterion for Extraintestinal pathogenic *E. coli* (ExPEC). Urinary pathogenic *E. coli* (UPEC) isolates were those strains which harboured ≥ 3 of virulence genes including, *yfcV*, *fyuA*, *vat* and *chuA*. The virulence factor (VF) score was the total number of virulence genes detected, adjusted for multiple detection of the *pap* operon (*30*).

### Antimicrobial susceptibility testing

The standard disk diffusion method on Mueller-Hinton agar was used to determine the antibiotic susceptibility patterns of 76 ST131 strains and results were interpreted according to the Clinical and Laboratory Standard Institute (CLSI) guidelines (*31*). The number of antibiotics to which the strain was resistant was considered as resistance score. Isolates with resistance to at least one representative of three or more antimicrobial classes were defined as multidrug resistant (MDR) (*32*). ESBL production was assayed using phenotypic combined disk test according to the recommendations of CLSI (*31*).

### Detection of resistance encoding genes

The presence of carbapenemase (*bla*_IMP_, *bla*_VIM_, *bla*_KPC_, *bla*_NDM_ and *bla*_OXA-48_) (*33*), ESBLs (*bla*_TEM_, *bla*_SHV_, *bla*_OXA-1_, *bla*_CTX-M-15_ and *bla*_CTX-M_ groups 1, 2, 8, 9, 25) (*34*), plasmid mediated quinolone resistance (PMQR) (*qnrA*, *qnrB*, *qnrS* and *aac-6Ib-cr*) was investigated by multiplex PCR according to previously published methods (*35*). Furthermore, isolates harbouring the 16S *rRNA* methylase genes (*ArmA*, *rmtB*, *rmtC*) and aminoglycoside resistance determinants (*aac3IIa*, *aac-6Ib*) were detected by single PCR (*36, 21*).

### Statistical analysis

In order to compare the proportions and scores, the Fisher’s exact test and the Mann-Whitney U-test were used, respectively. The principal coordinate analysis, a multidimensional scaling method analogous to principal component analysis, was used to collapse the molecular data set for simplified between group comparisons (*21*). Groups were compared on each of the first three coordinates, which captured most of the variance within the dataset using a two-tailed t-test.

## Acknowledgments

We thank Professor James R. Johnson for his invaluable commenting, and scientific editing. We also acknowledge his generous and continuing support and encouragement.

This work was supported fully by Semnan University of Medical Sciences (grant No. 730).

